# Population genomics of adaptive radiations: Exceptionally high levels of genetic diversity and recombination in an endemic spider from the Canary Islands

**DOI:** 10.1101/2024.05.13.593866

**Authors:** Paula Escuer, Sara Guirao-Rico, Miquel A. Arnedo, Alejandro Sánchez-Gracia, Julio Rozas

## Abstract

The spider genus *Dysdera* has undergone a remarkable diversification in the oceanic archipelago of the Canary Islands, ∼60 endemic species originated during the 20 million years since the origin of the archipelago. This evolutionary radiation has been accompanied by substantial dietary shifts, often characterized by phenotypic modifications encompassing morphological, metabolic and behavioral changes. Hence, these endemic spiders represent an excellent model for understanding the evolutionary drivers and to pinpoint the genomic determinants underlying adaptive radiations. Recently, we achieved the first chromosome-level genome assembly of one of the endemic species, *D. silvatica,* providing a high-quality reference sequence for evolutionary genomics studies. Here, we conducted a low-coverage based resequencing study of a natural population of *D. silvatica* from La Gomera island. Taking advantage of the new high-quality genome, we characterized genome-wide levels of nucleotide polymorphism, divergence, and linkage disequilibrium, and inferred the demographic history of this population. We also performed comprehensive genome-wide scans for recent positive selection. Our findings uncovered exceptionally high levels of nucleotide diversity and recombination in this geographically restricted endemic species, indicative of large historical effective population sizes. Furthermore, we identified genomic regions potentially under positive selection, shedding light on relevant biological processes, such as vision and nitrogen extraction as possible targets of adaptation and eventually, as drivers of the species diversification. This pioneering study in spiders endemic of an oceanic archipelago lays the groundwork for broader population genomics investigations aimed at understanding the genetic mechanisms driven adaptive radiations in island ecosystems.

## Introduction

Understanding how species originate stands as a central issue in evolutionary biology (Austin & Arnold, 2001; Ravinet et al., 2017). Yet, despite its pivotal role in biodiversity management and conservation in the face of climate change (Mergeay & Santamaria, 2012), our knowledge of the evolutionary mechanisms and the key genomic targets driving species diversification remains limited. Certainly, evolutionary inference in rapidly evolving traits is more feasible in organisms with shorter generation times like viruses or bacteria (Coyne & Orr, 2009). Island radiations, especially those occurring in oceanic archipelagos offer experimental conditions for studying diversification in animals and plants. The small size and clear boundaries of oceanic islands, jointly with their simplified ecosystems that are repeated in the same or different oceans (resulting in multiple independent replicates of evolution), make islands test tubes for evolutionary studies (Hodges & Derieg, 2009; Schluter, 2000). For this reason, the oceanic archipelagos have long been recognized as natural laboratories to study short-term evolution (Fernández-Mazuecos et al., 2020), having received much attention over time (Carson & Kaneshiro, 1976; Emerson, 2002; Gillespie, 2004; Grant & Grant, 2008; Juan et al., 2000; Machado et al., 2017). In recent years, the advent of whole-genome data analysis, especially in non-model organisms, have further fueled the interest on adaptive radiations (Choi et al., 2021; Feder et al., 2012; Lamichhaney et al., 2015; Richards et al., 2021; Schwager et al., 2017; Wolf & Ellegren, 2017). However, the relative contributions of adaptive and non-adaptive forces in species diversification are still a matter of debate (Choi et al., 2020; Muschick et al., 2012; Rundell & Price, 2009; Rundle & Nosil, 2005; Simões et al., 2016).

The Canary Islands, a volcanic archipelago comprising eight islands, exhibit significant climate variations (Fernández-Palacios, 2011) and high levels of endemism, particularly among Arthropods, reaching up to 40% (Martín et al., 2010). Among the endemic species, spiders of the genus *Dysdera* Latreille, 1804 (Araneae: Dysderidae) represent one of the most spectacular examples of islands radiation among spiders (Arnedo et al., 2001, 2007; Macías-Hernández et al., 2016; Řezáč et al., 2021; Bellvert et al., 2023). Indeed, about 20% of the ∼300 species described in this Western Palearctic genus (World Spider Catalog, 2024) are endemic to this archipelago. *Dysdera* spiders are nocturnal ground-dwelling hunters recognised as one of the few examples of prey specialization (stenophagy) among spiders (Pekár et al., 2016). Some species are facultatively or even obligatorily specialized in eating woodlice, a prey that most predators commonly avoided due to their effective curling defense strategy, and the accumulation of heavy metals in their exoskeleton, making them difficult to capture and metabolize (Pekár et al., 2016). These spiders have developed morphological adaptations, behavioral strategies and metabolic mechanisms to successfully capture and digest these isopods (Hopkin & Martin, 1985; Řezáč et al., 2008; Řezáč & Pekár, 2007; Toft & Macías-Hernández, 2017; Bellvert et al., 2023). Dietary diversification appears to be directly linked to species richness and species overlapping in the same locality (Řezáč et al., 2021). In a first attempt to determine the molecular basis of such extraordinary dietary adaptations, we carried out a comparative transcriptomic analysis involving species exhibiting differing levels of stenophagy from two evolutionary independent shifts towards feeding on woodlice (Vizueta et al., 2019). This study uncovered several candidate genes likely associated with diet diversification within this genus in the archipelago. Specifically, we identified genes involved in heavy metal detoxification and homeostasis, as well as in the metabolism of crucial nutrients and venom toxins. Recently, we reported a chromosome-scale genome assembly of *D. silvatica* (Escuer et al., 2022), a species with low levels of dietary specialization that inhabits the three more western Canary Islands (La Gomera, La Palma and El Hierro; (Macías-Hernández et al., 2016). The genome size of *D. silvatica* is estimated to be 1.7 Gb based on flow cytometry (1.4 Gb in the published assembly), organized into seven holocentric chromosomes (Schrader, 1935). Here, we utilized this high-quality reference genome to conduct the first population genomics study in a *Dysdera* species. Our main goal was to characterize and quantify the levels and patterns of natural genomic variation in an endemic species from the Macaronesia archipelago and assess the impact of recent selection on genome-wide patterns of polymorphism and divergence. While other population genomic studies in spiders have been published, very few have used whole-genome data (Chen et al., 2020; Hendrickx et al., 2022; Huang et al., 2022). We used a low-coverage sequencing strategy, which has proven to be a cost-effective approach for conducting population genomic studies, particularly in non-model species with large genome sizes and having a gold-standard reference genome (Lou et al., 2021). We have analyzed genome-wide patterns and levels of nucleotide variation, and linkage disequilibrium (LD), and estimated per site recombination rates in 12 genomes of *D. silvatica* sampled at a single locality in La Gomera. Our analyses, which include demographic and positive selection inference, reveal a set of candidate genes associated with key biological functions such as vision and nitrogen extraction. These genes emerge as potential targets of adaptation and may play a key role as drivers of species diversification. Our findings contribute to the understanding of one of the least studied groups of animals at the population genomic level, with demonstrated importance for ecosystems, productivity and health. Still, this group poses important methodological challenges due to their large genome and chromosome sizes and high repetitive content.

## Material and Methods

### Sampling, DNA extraction and whole-genome sequencing

We collected 12 individuals of *D. silvatica* (two males and 10 females) from the same locality in La Gomera Island (Teselinde; Ermita de Santa Clara, Vallehermoso; Supplementary Table S1). Additionally, we collected one male of *D. bandamae* from Gran Canaria (Llanos de la Pez; Tejada; Figure 1; Supplementary Table S1) to be used as the outgroup for evolutionary inference. For DNA extraction, we used a modification of the Gentra Puregene Cell kit (Qiagen) specifically optimized to enhance extraction efficiency in chelicerates. The extracted DNA was sequenced on the NovaSeq 6000 (for all *D. silvatica* samples) and HiSeqX (for the individual of *D. bandamae*) platforms in Macrogen Inc. (Seoul, Korea).

**Figure 1.**
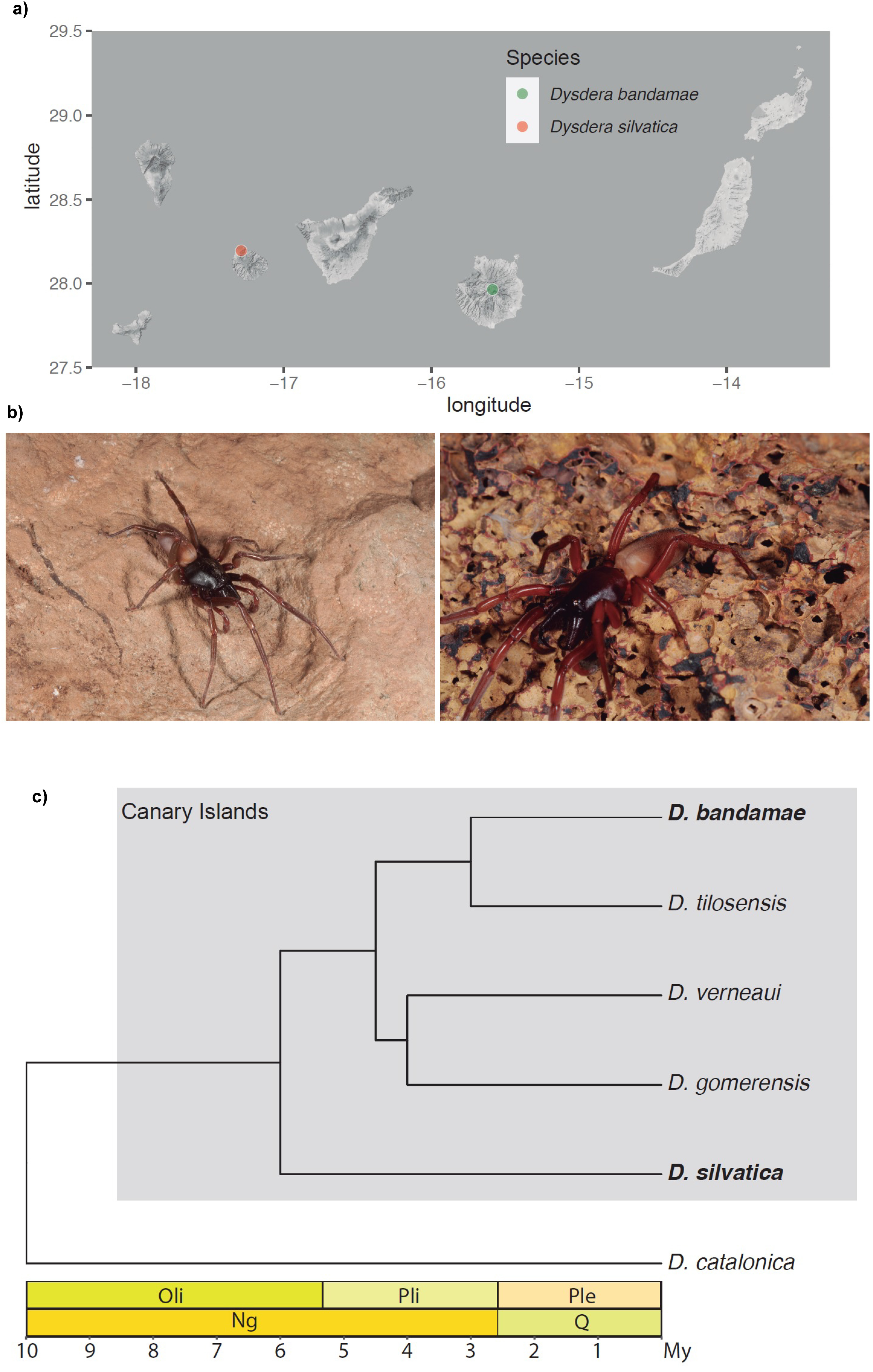
Distribution area, phenotypic traits and phylogenetic relationships of five *Dysdera* species from the Canary Islands. a) Map of Canary Islands showing the geographical localization of the species used in this project. b) Images of the studied *Dysdera* species; left, *D. silvatica*; right, *D. bandamae*. c) Phylogenetic relationships among some *Dysdera* species.

Due to the large genome size of *D. silvatica* (about 1.7 Gb estimated from flow cytometry; Sánchez-Herrero et al., 2019), we opted for a low coverage whole-genome resequencing strategy to generate the *D. silvatica* intraspecific data. We obtained an average raw sequencing depth for each *D. silvatica* individual of 6.7x (Supplementary Table S1), and 31.8x for *D. bandamae* (Supplementary Table S1). We also obtained the genome sequence of one female of *D. silvatica* at medium coverage (41.2x).

### Read quality assessment, trimming and mapping

We used FASTQC v0.11.9 (Andrews, 2010) to assess the quality of raw reads and Trimmomatic v0.39 (Bolger et al., 2014) for the trimming process. We filter out adapters, all reads shorter than 50 bp, or those with bad quality scores (< 15) across sliding windows of 4 bp length. We also removed leading and trailing bases with low quality scores (< 3) and all missing bases.

Filtered reads were mapped independently for each individual of both species, against the reference genome of *D. silvatica* (Escuer et al., 2022) using bwa mem v0.7.16 (Li & Durbin, 2009) with short split hits labeled as secondary alignments (-M). Then, we used Samtools v1.11 (Danecek et al., 2021) to filter out these secondary alignments. We removed duplicates from the resulting BAM files, added read groups labels, and the alignments were indexed and sorted using Picard Tools v2.26.10 (Broad Institute, 2016). Finally, we estimated the average and median read depth for each scaffold using Samtools v1.11 and Bedtools v2.29.1 (Quinlan & Hall, 2010), along with Qualimap v2.2.1 (Okonechnikov et al., 2016). The average net coverage per site of the *D. silvatica* sample is 44.3x, with the average net coverage per individual ranging from 3.3x to 4.3x (Supplementary Table S2). Since *D. silvatica* males are X0, the average sequencing depth in the X chromosome is about half that of autosomes (Supplementary Table S2). We conducted all analysis separately for the autosomes (*n* = 24) and the X chromosomes (*n* = 20). We excluded the two males from the X-chromosome analysis since ANGSD (Korneliussen et al., 2014) cannot handle haploid data (i.e., it calculates diploid genotype likelihoods). The average net coverage of the female sequenced at medium coverage is 33.1x.

### Genome-wide nucleotide polymorphism and divergence

We utilized ANGSD along with related software (e. g. ngsLD (Fox et al., 2019), ngsDist (Vieira et al., 2016) and ngsTools (Fumagalli et al., 2014), which consider genotype uncertainty for the downstream population genomic analysis of the low-coverage samples.

To obtain a consensus sequence of *D. bandamae* in FASTA format we use the `-doFasta` option of ANGSD. Before estimating GLs, we applied specific quality filters in ANGSD to remove non-informative positions. We filtered out positions with very low or very high net coverage across samples (`-setMinDepth 3, -setMaxDepth 92`), corresponding to the 2.5th and 97.5th percentiles of the per-site net coverage distribution across all individuals. In addition, we only retained positions with high base calling quality, properly paired and high mapping quality reads (`-minQ 20, -only_proper_pairs and -minMapQ 20`). We adjusted the mapQ parameter for excessive mismatches (`-C 50`) and the qscores around indels (`-baq 1`), and discarded not primary, failure and duplicate reads (-remove_bads), as well as reads that did not map uniquely (`-uniqueOnly`). We set the *D. silvatica* genome as a reference genome (`-ref`) and estimated the GLs for each individual using the GATK algorithm (`-GL 2`). To mitigate potential bias introduced by repetitive elements, which might affect the mapping results, we conducted additional analyses with different coverage filters (`- setMinDepth 20, -setMaxDepth 72`), encompassing the 66.6% of the net coverage distribution, and excluding masked regions of the genome identified by RepeatMasker.

We analyzed the levels and patterns of nucleotide variation using the unfolded site allele frequency (SAF) likelihood obtained from the *realSFS* program within ANGSD and utilizing the sequence of *D. bandamae* as the ancestral reference (`-doMajorMinor 5, -anc`). We estimated the SAF based on individual GL assuming HWE (`-doSaf 1`). From the estimated SAF, and using the *thetaStat* program within ANGSD, we calculated several summary statistics and neutrality tests, including the number of polymorphic sites (*S*), nucleotide diversity (*π;*Nei, 1987), Watterson estimator of theta (*θ*_w;_ Watterson, 1975), Tajima’s *D* (Tajima, 1989), Fu and Li’s *D* and *F* (Fu & Li, 1993), Fay and Wu’s *H* (Fay & Wu, 2000) and Zeng’s *E* (Zeng et al., 2006). We computed these summary statistics and neutrality tests for both the entire chromosome and in a sliding window approach (in non-overlapping windows of 1 kb and 50 kb).

We used ngsDist v.1.0.10 (Vieira et al., 2016) to compute the nucleotide divergence between *D. silvatica* and *D. bandamae* based on the estimated GLs. For that, we first generated a GLs file in BEAGLE format using the BAM files of the 13 individuals (12 *D. silvatica* and one *D. bandama*e for the autosomes; 10 *D. silvatica* and one *D. bandamae* for the X chromosome). Using these GLs, we calculated the evolutionary distance between the two species as the average pairwise JC69 corrected distances between each individual of *D. silvatica* and *D. bandamae*`(--evol_model 2`; Jukes & Cantor, 1969). We computed nucleotide divergence for the entire chromosome and in non-overlapping windows of 1, and 50 kb).

We used Picard (Broad Institute, 2016) to incorporate read groups to the BAM file of the individual sequenced at medium coverage and called their variants with GATK HaplotypeCaller (Van der Auwera, & O’Connor, 2020). We selected SNP variants with total depth between 10 and 60, phred-scaled quality score > 30, normalized variant quality > 2, StrandOddsRatio < 3 and FisherStrand < 60 (both measuring phred-scaled probability of strand bias), and Mapping quality > 40, to generate the VCF of this individual using GATK VariantFiltration tool.

### Population structure and demography

We conducted a Principal Component Analysis (PCA) to explore population structure in our low-coverage sample. We used the ngsCovar software from ngsTools v3 (Fumagalli et al., 2014). This analysis was based on the estimated GL and was performed separately for autosomes and the X chromosome. We applied the same filters and options used to estimate GL for variation analyses but generating a binary file with GL (`-doGeno 32)`, assuming that we know major and minor alleles `(-doMaf 1)`, excluding triallelic positions (`- SkipTrialletic 1`), and filtering out those SNPs with *P*-values > 1 x 10^-6^ (`-snp_pval`).

We used Stairway Plot2 (Liu & Fu, 2020) and the maximum likelihood estimate of the unfolded SFS to infer the recent demographic history of the *D. silvatica* population. The analysis was conducted separately for autosomes and the X chromosome. We set the generation time to 1.5 years (Cooke, 1965), and the neutral mutation rate per-site and per-generation (*μ*) to 6.07 x 10^-9^. This mutation rate was obtained from the nucleotide divergence estimated between *D. silvatica* and *D. bandamae* (*K* = 0.120; Table 1), and the estimated divergence time for these species of 14.8 mya (Crespo et al., 2021).

**Table 1.**
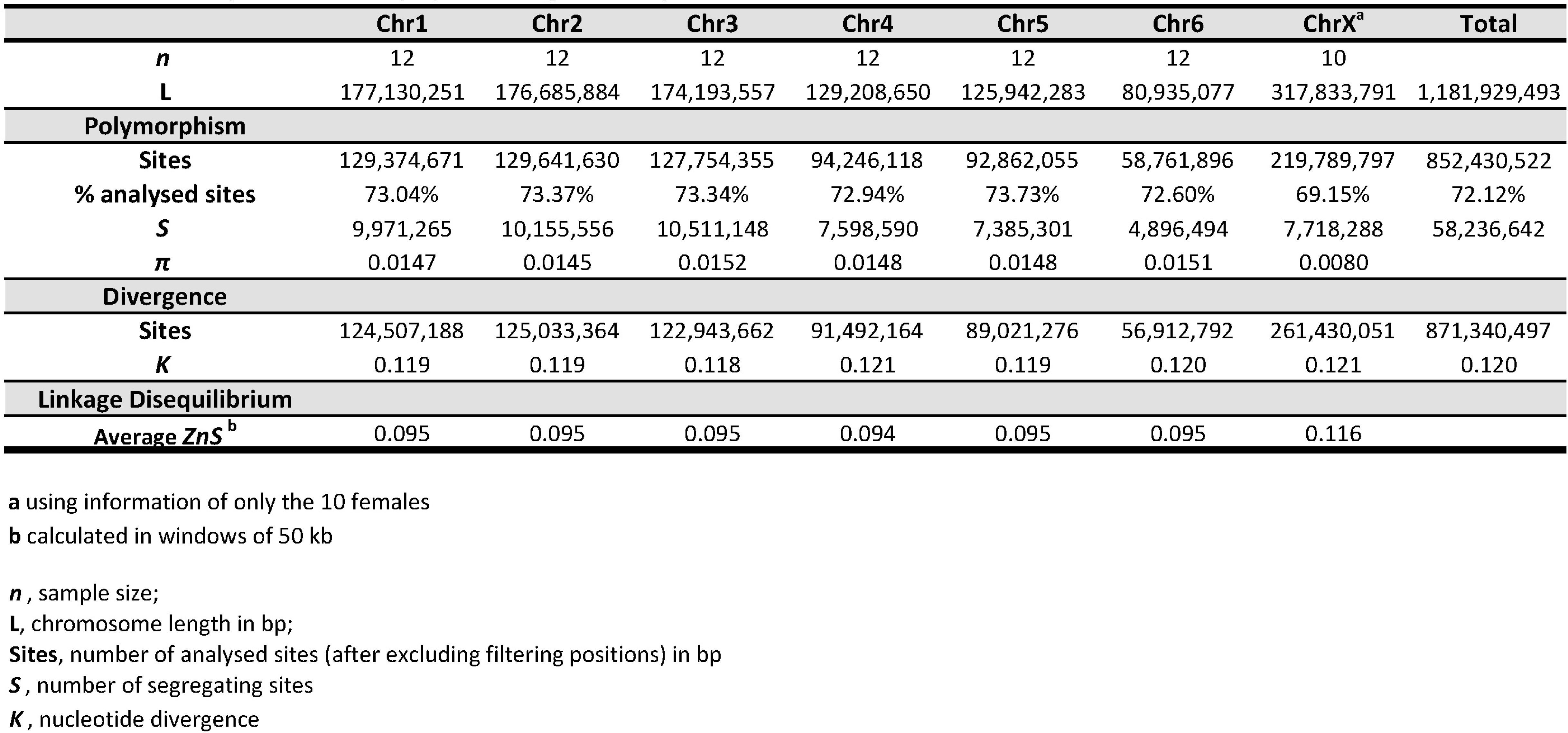
Summary statistics of population genomic parameters.

### Linkage disequilibrium and recombination rate

We estimated the site pairwise linkage disequilibrium (LD) using the ngsLD v1.1.1 software (Fox et al., 2019), taking genotype uncertainty into account. Initially, we generated a file with GL in BEAGLE format (`-doGlf 2`) with ANGSD by applying the same filters and options as described above. Then, LD was estimated as the average *r^2^* between genotypes (i.e., the *ZnS* statistic; (Kelly, 1997) in non-overlapping windows of 1.5 and 50 kb. To reduce the number of pairwise comparisons, we randomly sampled 0.1% of the sites to estimate the decay of LD with distance.

We used the composite-likelihood approach implemented in pyrho v0.1.6 (Spence & Song, 2019) to estimate fine-scale per-base per-generation recombination rate (*r*) in *D. silvatica* based on LD patterns. For each chromosome, we run pyrho under two scenarios: i) a constant size demographic model, using the size estimated for the present in the demographic analysis, and ii) the best-fit demographic scenario from this analysis. We used a multi-variant call format (VCF) file with SNPs called in ANGSD (`-doGeno 1, -doMajorMinor 5, -doPost 1, -skipTriallelic 1, -snp_pval 1e-6` and `-dobcf)` as the input for pyrho. We masked repetitive regions in the reference before the variant calling. The specific parameters set in pyrho can be found in Supplementary Table S3. We also used the sequentially Markovian coalescent model implemented in iSMC (Barroso et al., 2019) and the VCF of the individual of *D. silvatica* sequenced at medium coverage, to infer the population scale recombination rate (Rhro) in this species.

### Genome scans for selection

We used the RAiSD v2.9 software (Alachiotis & Pavlidis, 2018) to search for the characteristic hallmarks of positive selection within the *D. silvatica* genome. This method relies on a composite statistic (*μ-R*), calculated through a SNP-driven, sliding-window algorithm (using the default window length of 50 SNPs), integrating evidence indicative of selective sweeps, such as *μ-var* (reduction of nucleotide variation), μ*-sfs* (shifts to low and high-frequency derived variants), *μ-ld* (elevated levels of LD around the selective site). RAiSD uses the VCF file with called genotypes (see above) as the input. Subsequently, we utilized the script *vcfutils.pl* from Samtools v1.11 (Danecek et al., 2021) to convert the resulting BCF file into VCF format. We used the -X the option to exclude repetitive regions from the analysis.

We used a three-step approach to identify outlier genomic regions within the empirical distribution of *μ-R*. First, we retained windows exhibiting the top 0.1% values of the *μ-R* statistic. Second, we examined the retained windows for consecutive regions showing *μ-R* values above 0.01% of the genome-wide empirical distribution. Among these, we selected those with a minimum of two consecutive windows with *μ-R* values above 0.001%. We then used structural annotations within the reference genome to identify genes or other functional elements located within these outlier regions.

We also applied the McDonald and Kreitman test (MKT) (McDonald & Kreitman, 1991) to coding sequences to detect the footprint of positive and negative selection in the genome of *D. silvatica*. We first obtained a VCF file containing the genotypes of the 12 individuals of *D. silvatica* and the individual of *D. bandamae* using ANGSD. We applied the same filters and options as those used for estimating GL and for preparing the VCF input for pyrho. We used snpEff v.5.1d (Cingolani et al., 2012) to predict the functional effects of genomic variants (including synonymous and nonsynonymous mutations) across the VCF containing data from 13 individuals. The annotated VCF file was used to run the MKT on all functionally annotated protein-coding genes by using a modification of Tomas Blankers’ script (available at https://github.com/thomasblankers/popgen/blob/master/MKTtest). We filtered out sites with a minor allele frequency < 0.1 (i.e., singletons).

### Mitochondrial assembly and phylogenetic analyses

We assembled the mitochondrial genomes of *D. silvatica* (one per individual) and *D. bandamae* using NOVOplasty v4.3 (Dierckxsens et al., 2017) and the raw reads of these species. For the assembly, we applied a Genome Range of 14000-20000 bp and a K-mer size of 33. Subsequently, we generated a multiple sequence alignment of the genomes (13 individuals) using MAFFT (Katoh & Standley, 2016) with the following options `--maxiterate 1000, --globalpair` and the *G-INS-i* algorithm. We then built a phylogenetic tree with IQ-TREE2 v1.6.12 (Minh et al., 2020) with parameters `-m MFP -B 1000`, and visualized and edited the tree with iTOL web interface (Letunic & Bork, 2021).

### GO enrichment

We used InterProScan v.5.57-90.0 (Jones et al., 2014) to extract the GO terms associated with the genes annotated in the reference genome v.2.3 of *D. silvatica* (Escuer et al., 2022). For the GO enrichment analysis we used the R packages GSEABase v.1.60.0 (Geistlinger et al., 2021; Morgan et al., 2023), GOstats v.2.64.0 (Falcon & Gentleman, 2007) and org.Dm.eg.db v.3.16.0 (Carlson, 2019) with the GO associated to the protein-coding genes located in the outlier regions in the RAiSD analysis (Alachiotis & Pavlidis, 2018). We used the R package q-value v.2.30.0 (Storey, Bass, Dabney, 2022) to transform the obtained *P*-values into *q*-values. The GOstats analysis was run under the conditional option. We considered as significantly overrepresented those GO terms with an associated P-value < 0.01 or with q-values < 0.10. We used the R packages clusterProfiler v.4.8.1 (Wu et al., 2021) and AnnotationForge v.1.42.0 (Carlson & Pagès 2023) to transform the obtained results from GOstats into an enrichResult object and plotted the results employing the R packages enrichplot v.1.20.0 (Yu, 2023), europepmc v.0.4.1 (Ferguson et al., 2021) and ggplot2 v.3.4.2 (Wickham, 2016). The library enrichplot was used to reduce the complexity of the enriched GO terms for better interpretability of the results. The number of clusters (nCluster option) was set to the default value of 5.

## Results

### High levels of nucleotide polymorphism and recombination in *D. silvatica*

After excluding positions that did not meet strict quality filters, we analyzed 632.64 Mbp and 219.80 Mbp in autosomes and the X chromosome, respectively, which correspond approximately 72% of the total sequenced genomic positions (Table 1). These positions contained 50,518,353 autosomal and 7,718,288 X chromosome SNPs. Levels of nucleotide diversity are notably high in *D. silvatica* autosomes (*π* = 0.015; Table 1; Supplementary Tables S4-S5) and evenly distributed across the genome (Table 1; Figure 2a). As expected, the X chromosome is less variable, owing its smaller effective size (*π* = 0.008; Table 1; Figure 2a). Tajima’s *D* values are consistently negative across the genome, although with values not reaching extreme levels (Table 2; Supplementary Tables S4-S5). Overall, autosomes tend to have more negative Tajima’s *D* values than the X chromosome (Table 2; Supplementary Tables S4-S5). Nucleotide divergence is also homogeneously distributed across chromosomes, with an average *K* value of 0.120 (Table 1; Figure 2b).

**Figure 2.**
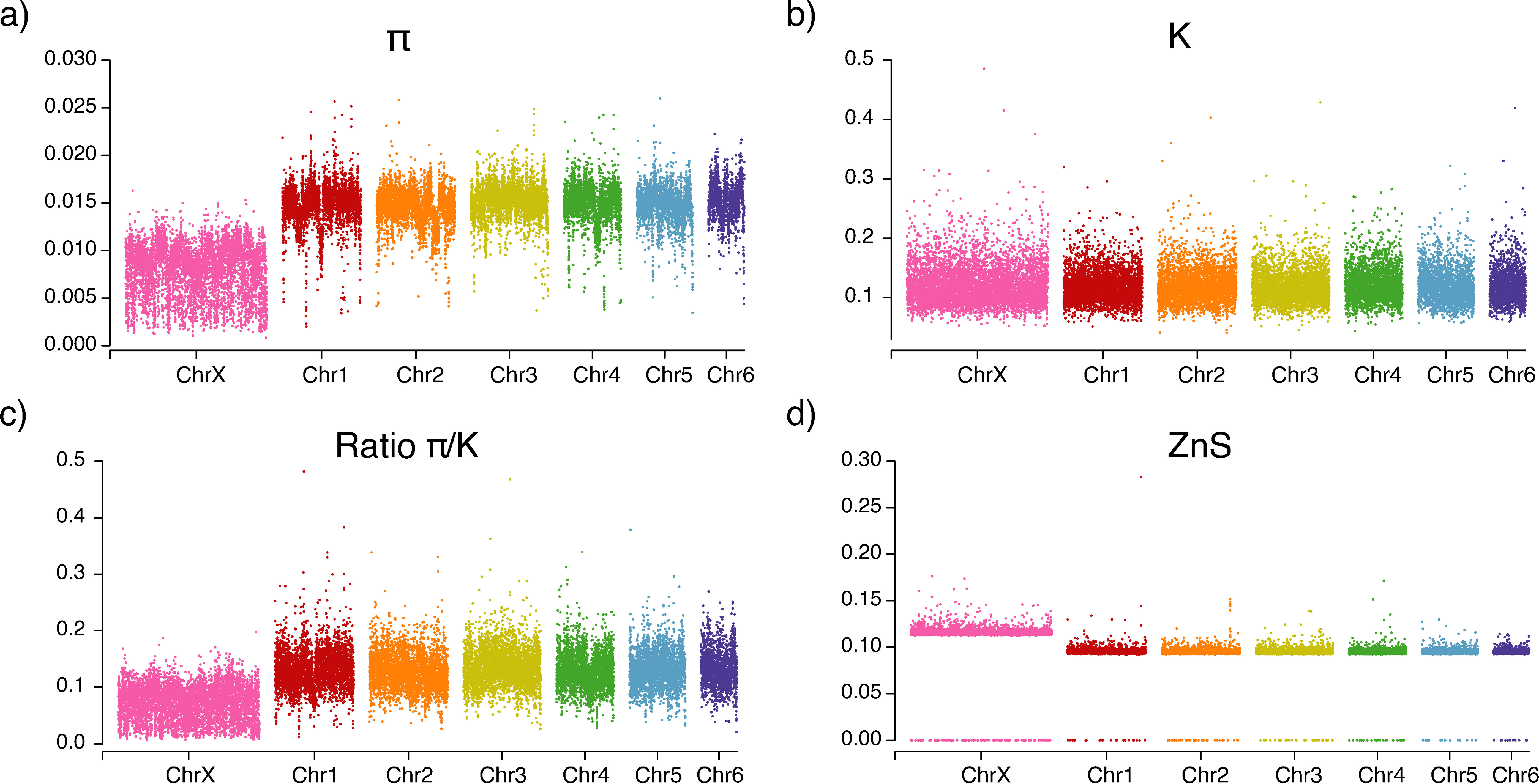
Genome-wide distribution of summary statistics of genome variation. a) Nucleotide diversity (*π*); b) Nucleotide divergence per site (*K*); c) Ratio *π/K*; d) Linkage disequilibrium (estimated as ZnS). Each point depicts the value calculated in a window of 50 kb.

**Table 2.**
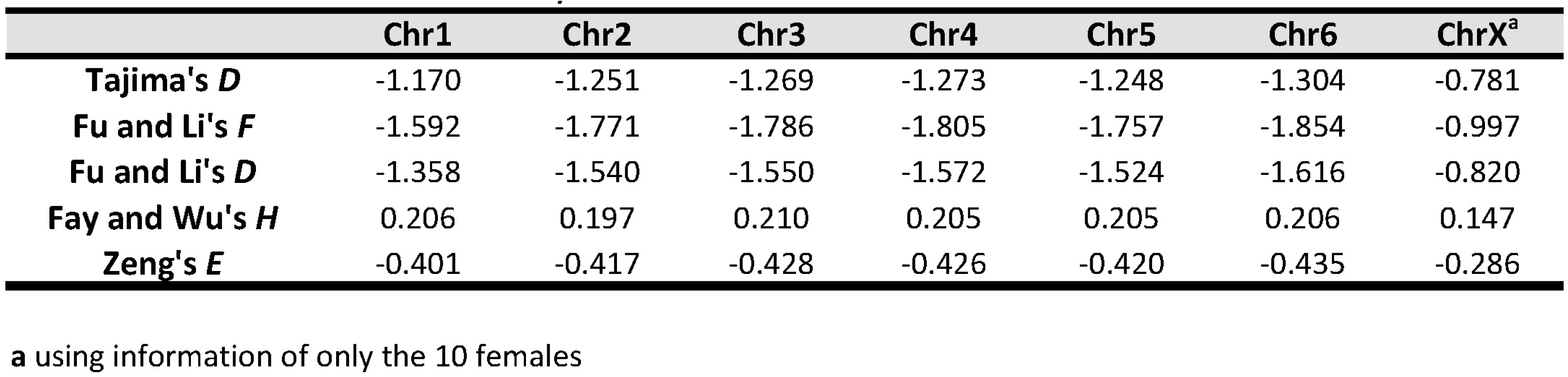
Results of the neutrality tests.

Linkage disequilibrium, estimated as *ZnS*, exhibits a rapid decay with distance (Table 1; Figure 2d, Supplementary Table S6), suggesting high levels of genetic recombination in the genome of this species. As expected, *ZnS* values are consistently higher in the X chromosome compared to the autosomes (Table 1; Figure 2d). In fact, the estimates of local per-base per-generation recombination rate (*r*) is unusually high in this species, regardless of the underlying demographic model used for the inference (see below) or the chromosome analyzed, with most genomic windows displaying values of *r* between 10^-7^ and 10^-6^ (Supplementary Table S3). These estimates are generally higher in autosomes than in the X chromosome (Supplementary Tables S3). We found significant but modest correlation between *r* and *π*, with X the chromosome exhibiting higher correlation coefficients than autosomes (Supplementary Figures S2 and S3).

### Large historical effective population sizes, recent decline and no evidence of population structure

Autosomes and the X chromosomes show concordant demographic histories (Figure 4), with the expected differences in parameter estimates resulting from the smaller effective population size of the latter. We inferred two major demographic events in the recent history of *D. silvatica* populations that are supported by bootstrap analysis. The older event reflects a population bottleneck that occurred around 200 kya, resulting in a five-fold reduction in population size, which persisted for ∼35 kya. Following the complete recovery, the effective size of the population remained stable until around 10 kya, when an abrupt reduction dropped the population size to *N*_e_ of ∼25 x 10^3^ individuals.

**Figure 3.**
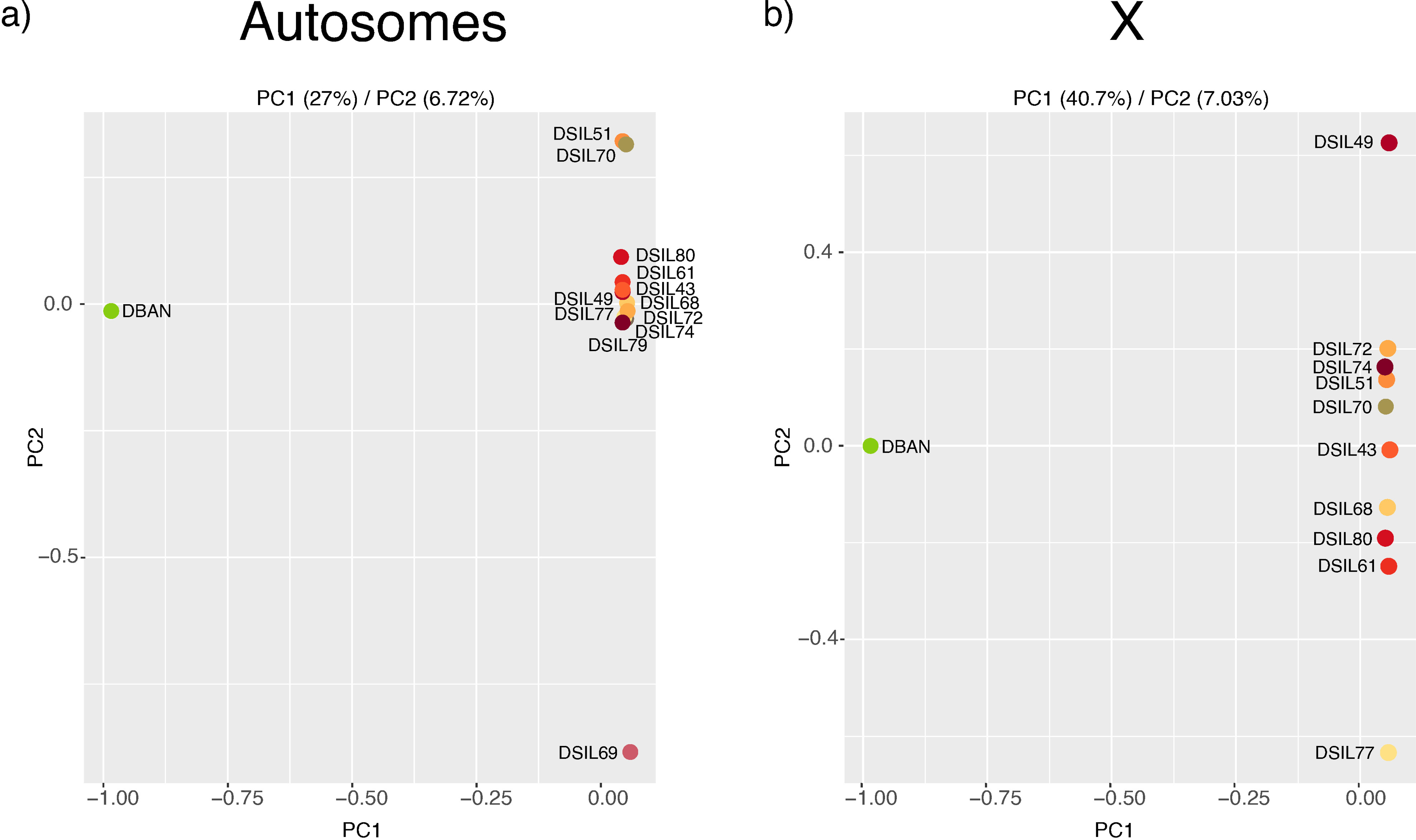
Principal Component Analysis (PCA) using data from a) autosomes and b) X-chromosomes (data from only the 10 females).

**Figure 4.**
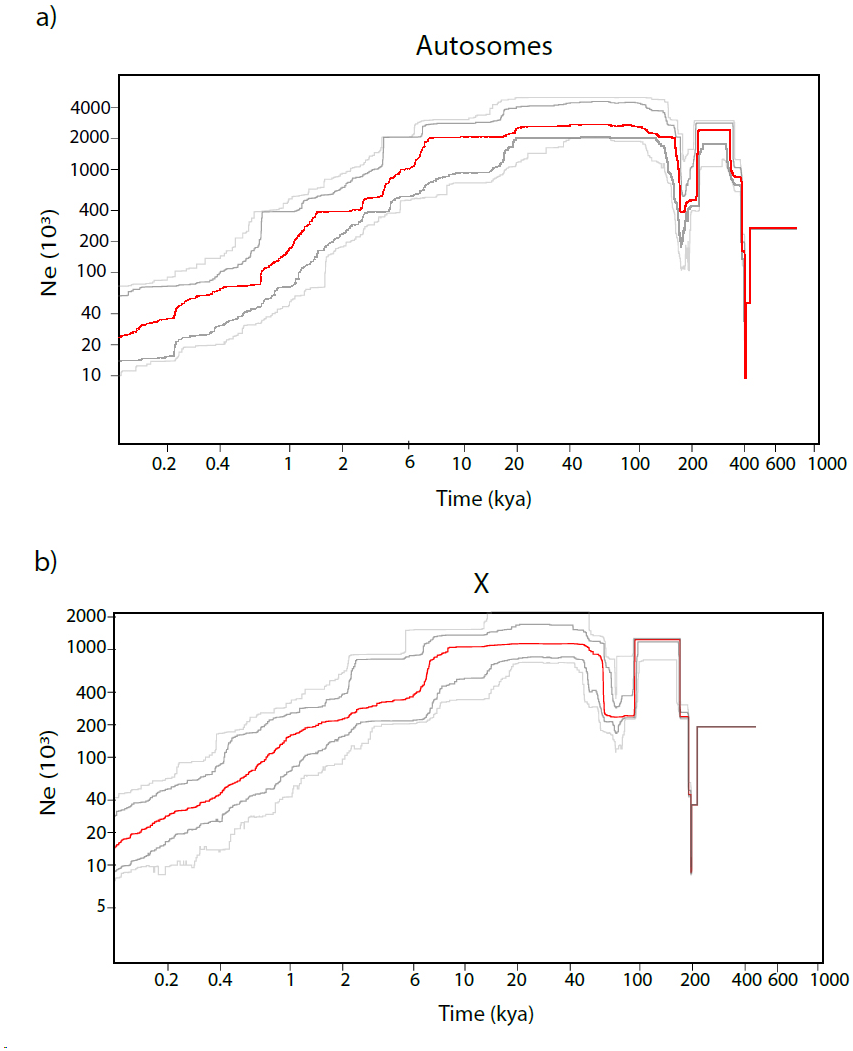
Population demographic history inferred from a) autosomal data (*n* = 24), and b) X-chromosomes (*n* = 20). The red line indicates the median population size (200 bootstrap replicates). Dark-gray and light gray lines delimit the 75% and the 95% bootstrap confidence intervals, respectively. Inference based on the unfolded SFS. Mutation rate per site and per generation of 6.07 x 10^-9^. Generation time of 1.5 years.

On the other hand, no evidence of population structure was found in the surveyed population (Figure 3; Supplementary Figure S1). While PCA results suggest some degree of differentiation between some *D. silvatica* individuals with respect to PC2 (Figure 3), this component only explains 7% of the variance, and the individuals involved are different in the different chromosomes. As expected, most of the variance explained by PC1 is attributed to the divergence between species. Similarly, there are no apparent signs of structuring in the phylogenetic tree based on mitogenomes (Supplementary Figure S1).

### Natural selection in recent past of *D. silvatica*

We identified 17 genomic regions as candidates to have been the target of positive selection in *D. silvatica*. Among them, 10 regions exhibit two or more consecutive windows with *μ-R* values above the 0.001% threshold of the empirical distribution (Supplementary Tables S7 and S8). Notably, these regions are overrepresented in the X chromosome, comprising 52% of candidate regions despite accounting for only 26% of the genome assembly (chi-square test *P*-value < 10^-5^). The windows with highest *μ-R* statistic values, as well as the largest number of consecutive significant windows, are predominantly found in chromosome 1 (Figure 5). In this chromosome, we identified a particularly strong candidate region (designated as Chr1_ID1) spanning over 146 kb with a maximum *μ-R* value of 228 (Supplementary Table S8).

**Figure 5.**
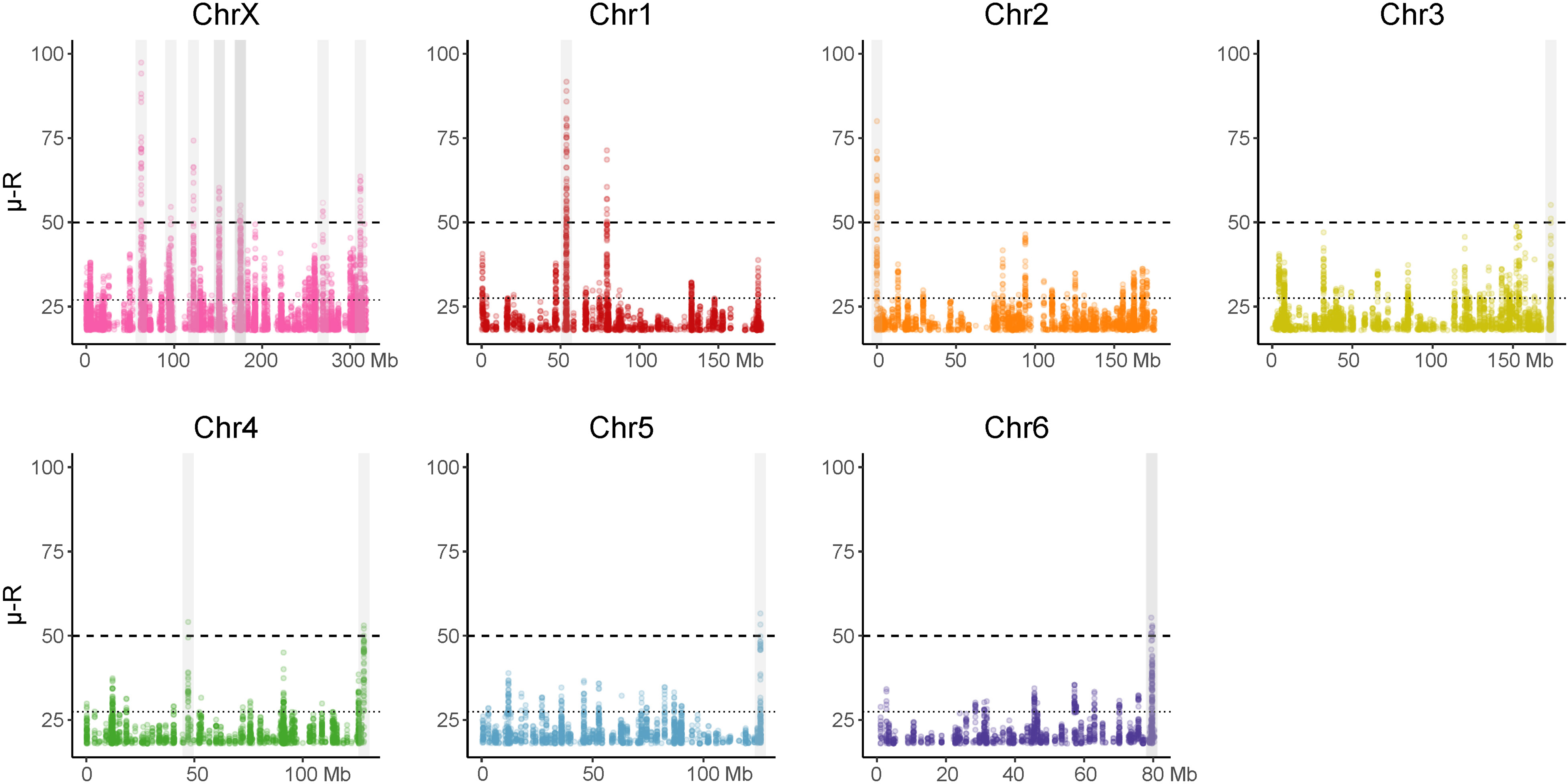
Results of the genome scan of selection. Dots depict those values of the *μ-R* statistic. Dotted and dashed lines indicate the cut-off values at 0.0001% and 0.00001%, respectively (Supplementary Table S7). Outlier regions analyzed are shaded in gray.

We identified 46 protein-coding genes annotated within the 17 candidate regions for positive selection. The GO enrichment analysis on the terms associated with these 46 genes revealed significant enrichment across 26 biological processes (BP), 14 molecular functions (MF), and 1 cellular component (CC) (*P*-value < 0.01; Table S9). However, after controlling for the false discovery rate (FDR), only 3 BPs and 14 MFs terms remained significant (*q*-value < 0.10; Table S9). The enriched biological processes are associated with genes involved in the visual system, nutrient transport, biosynthetic processes and apoptosis, among others. We identified molecular functions related to glycerol, sucrose and glutamate. However, none of these functions remained significant after correction for FDR (*q*-value > 0.05; Table S9B). We also found one significantly enriched cellular component, which is associated with the helicase complex, although with *q*-values above the significance threshold (Table S9C). Notably, only GO terms associated with the visual system and apoptosis remain significant after applying the correction for multiple testing (Table S9).

To gain further insights into the biological relevance of the aforementioned results, we also performed a hierarchical clustering of the enriched GO terms across the three GO domains, a method based on a similarity index between GO terms. The names of the main clusters and the number of GO terms within each cluster varied slightly depending on whether the *P*-value (Figure 6) or the *q*-value (Figure 7) was used as the threshold to select enriched terms. However, in all cases, the main clusters of enriched GO terms are related to the regulation of photoreceptor development, biosynthesis of components of the arginine pathway, chitin biosynthesis, glycerol catabolism, and sucrose transport. The GO terms associated with the visual system consistently display the lowest *P-* and *q*-values, and are among those with the highest number of associated genes.

**Figure 6.**
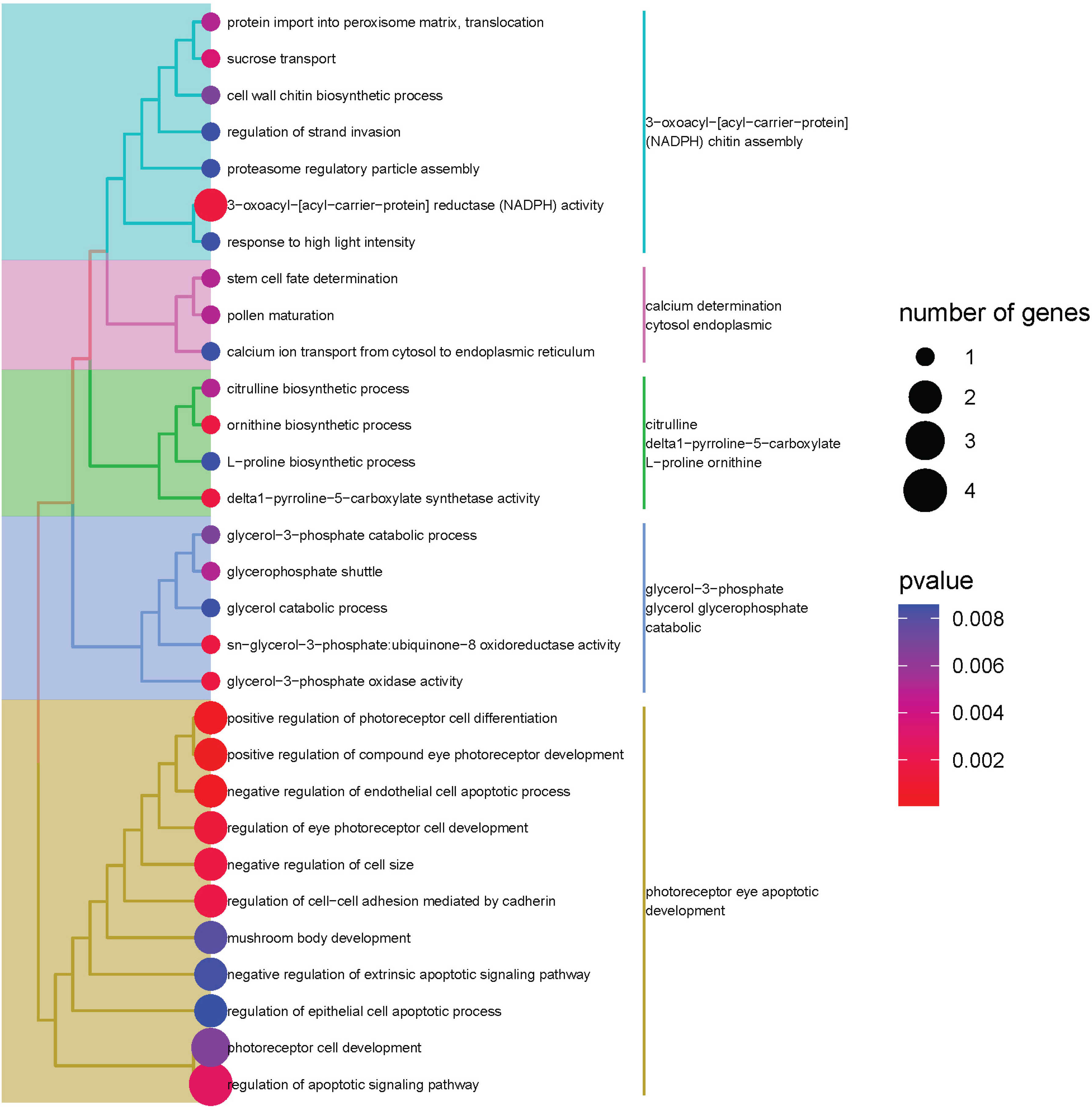
Hierarchical clustering of significantly enriched terms (*P*-value < 0.01). Clusters are highlighted in different colors.

**Figure 7.**
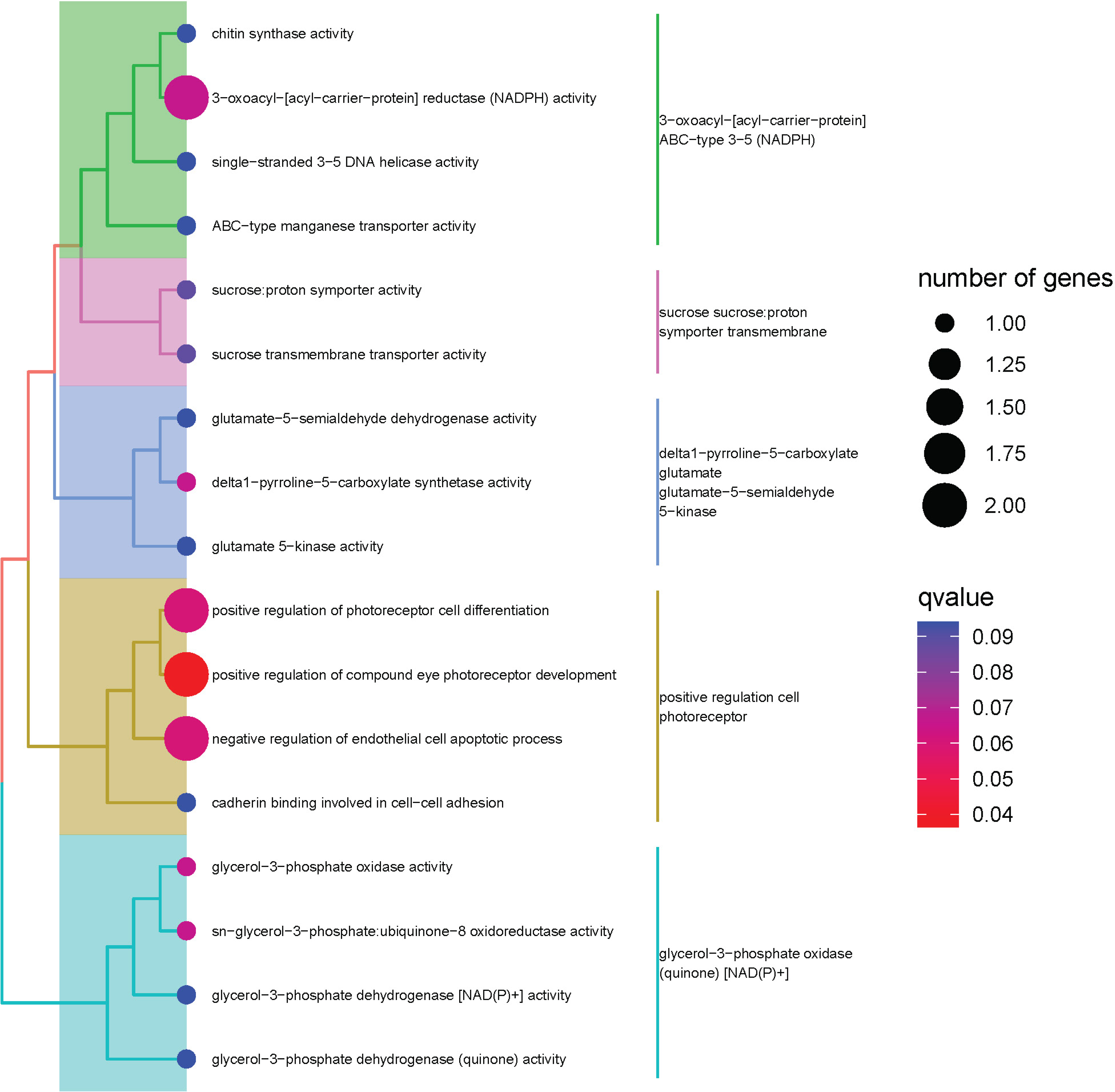
Hierarchical clustering of significantly enriched terms after correcting by FDR. Clusters are highlighted in different colors.

We applied the MKT to 28,900 genes (i.e., all putatively functional protein-coding genes annotated in the *D. silvatica* reference genome). Unfortunately, the vast majority of them (25,657) lacked sufficient divergence to obtain reliable results in this neutrality test. Among the 3,243 analyzed genes, 104 showed statistically significant departures from neutral expectations (*P*-value < 0.05), with 49 and 55 of them showing signals of negative or positive selection, respectively (Supplementary Table S10).

None of the genes exhibiting footprints of natural selection in the divergence of *D. silvatica* and *D. bandamae* are within the candidate regions identified by the RAiSD analysis, suggesting that the targets of selection are different at different evolutionary time scales. However, we found enriched GO terms associated with the visual system that are shared between the two analyses (Supplementary Table S11). This indicates that the visual system has been recurrently targeted for adaptation throughout the recent history of this species, albeit mediated by different target genes.

## Discussion

In this study, we present the first comprehensive population genomics analysis of an endemic spider from the Canary Islands, the nocturnal ground-dwelling spider *D. silvatica*. We adopted a low-coverage whole-genome resequencing strategy since it represents an optimal balance between inference power and sequencing costs, particularly suited for organisms with large genome sizes. Surprisingly, despite being an endemic species confined to three of the smallest islands of the Canarian archipelago, our analysis revealed that *D. silvatica* harbors exceptionally high nucleotide diversity levels (*π* = 0.015; for autosomes; Table 1), which approach the median diversity observed in Arthropods (Romiguier et al., 2014).

The scarcity of population genomics studies in island endemic species, other spiders, or even other chelicerates challenges the interpretation of the high levels of polymorphism detected. Our estimates for *D. silvatica* are much higher than those calculated in the few available studies within chelicerates, namely mites (*π =* 0.0006; Chen et al., 2020) and social spiders (e.g., *Stegodyphus sarasinorum*; *π =* 0.0005; Settepani et al., 2017). Indeed, these values even exceed genome-wide averages reported for the cosmopolitan fruit-fly *Drosophila melanogaster* (averages of *π* = 0.003-0.006) and overcome those estimated in high-recombination regions in this species (*π* < 0.01; (Kapun et al., 2021). Under the mutation-drift equilibrium, these elevated levels of nucleotide polymorphism would indicate either large historical effective population sizes or high mutation rates in this species. Based on divergence data, we estimated the neutral mutation rate per site and per generation to be *μ* = 6.07 x 10^-9^, which is a value typically observed in eukaryotes. Then, assuming a random mating, constant size population, the estimated nucleotide diversity values would imply a large effective population size (E(*N*_e_)= 375,000 individuals). Indeed, linkage disequilibrium and recombination rate estimates seem to anticipate even higher *N*_e_ in the past history of this species (see below).

The levels of intraspecific variation, but not the divergence estimates, differ between the X chromosome and the autosomes of *D. silvatica*, likely reflecting the expected smaller effective size of the sex chromosome. Even so, the X/autosomes *π* ratio (0.53) is lower than theoretically expected (0.75) and even far from the median value estimated for Arthropods (∼1; Leffler et al., 2012), indicating that other mutational or selective factors may be reducing the variability in the X chromosome. Moreover, our results do not support the ‘faster X’ hypothesis (Bechsgaard et al., 2019; Charlesworth et al., 2018) since the number of protein-coding genes with evidence of positive selection inferred by the MKT is not proportionally higher on the X than on the autosomes (Supplementary Table S10). However, the number of genes and substitutions used in the MKT analysis is too low to draw a firm conclusion. Data from a more distant species will be necessary to extend the analysis to the rest of the genes.

Results from demographic inference based on the unfolded SFS predict large ancestral population sizes for *D. silvatica* (∼2 x 10^6^ individuals). Despite the recent, abrupt decline in population size, we still observe remarkable levels of nucleotide diversity in this species. In the absence of clear signals of structure in both nuclear chromosomes and mitogenomes, we can rule out a recent admixture of highly diverged populations as the cause of these high levels of variability. Moreover, the levels of polymorphism estimated in the single individual sequenced at medium coverage (mean π = 0.016 for autosomes) are congruent with those estimated for the whole sample, indicating that the fine structure observed in the PCA does not account for this large, unexpected levels of polymorphism. Hence, this species appears to have maintained large, constant population sizes for a long time, at least during the last 100,000 years. The small proportion of nucleotide substitutions observed in coding regions between *D. silvatica* and *D. bandamae* suggest that this demographic history may be common to other, closely related species, pointing to shared polymorphism as an important factor to consider in variability studies on this genus in the Canary Islands.

The recombination rate also appears to be noticeably high across the *D. silvatica* genome, with an LD decay at the same order as that found in species with known large population sizes (Signor et al., 2018). Inaccuracies in the estimated demographic model and/or the scaling applied internally in pyrho to convert the ⍴ to *r*, not yet validated in non-human genomic data, may influence the magnitude of the estimated *r*. However, direct estimates of ⍴ in iSMC (per 100kb window autosomal ⍴ = 0.187), a method that jointly estimates demography and recombination, are consistent with a high recombination scenario. In any case, the estimated recombination landscape across the genome is expected to be confidently inferred. In this context, autosome recombination rates and LD are higher than those for the X chromosome (Figure 2d). Intriguingly, we have not detected visibly reduced levels of genetic recombination in the extremes of chromosomes, such as those observed in centromeric and telomeric regions in *Drosophila* (Comeron et al., 2012; Supplementary Figure S2). In line with this finding, we also failed to observe any obvious reduction in the variability of these chromosomal regions (Figure 2a). These findings may be connected with this species’ unique organization of holocentric chromosomes (Benavente & Wettstein, 1980). Indeed, the presence of holocentromeres does not appear to cause a reduction in the rate of recombination. In fact, in some cases, these levels are even remarkably high (e.g. Lepidoptera, Nematoda, Carex; (Escudero et al., 2018; Rockman & Kruglyak, 2009; Torres et al., 2023). Investigation into the role of these chromosomal organizations in shaping genomic features deserves attention in future studies.

In this study, we have identified several genomic regions and protein-coding genes with signals of recent positive selection in *D. silvatica* or after the split of this species and *D. bandamae*. Interestingly, several GO terms enriched in these features are of biological significance for this species. It is important to clarify here that we included in the analysis all GO terms, including those with few associated genes. The main reasons for not filtering out these terms are that i) we are dealing with a small list of candidate genes, and ii) we are studying a non-model-organism, for which we lack precise information on the function of genes in any closely related species; ontology information is therefore primarily based on orthology relationships with distant species. Overall, these two aspects are expected to affect statistical power in the GO overrepresentation analysis negatively. Overall, we identified a correspondence between the number of genes per GO term in our analysis and those found in former studies (published between 2010 and 2024) that include the same or related terms (Supplementary Figures S4 and S5). It is worth noting that those significantly enriched terms with a higher number of genes are those within the “photoreceptor” cluster (the cluster with lowest *P*- and *q*-values). These terms are among the ones having more functional information in databases, which makes us confident that vision is the biological process with best evidence of being the target of selection in *D. silvatica*. *D. silvatica* is a nocturnal ground*-*dwelling hunting spider that shelters in silk cocoons under rocks, tree trunks and barks during daylight (Pekár et al., 2016). Unlike many spiders, *Dysdera* lacks the anterior median eyes (AME, principal eyes), which are anatomically and functionally different from the remaining three pairs of eyes (secondary) (Morehouse et al., 2017). Principal eyes are responsible for spatial acuity and, in some spiders, color vision, whereas secondary eyes have less acuity and specialize in peripheral view and movement detection. In many cases, secondary eyes possess a mirror -like tapetum, a biological reflector enabling nocturnal organisms to see in dim light. In visually guided animals, one of the evolutionary strategies to enhance the visual system is by increasing eye size (Gonzalez-bellido et al., 2022). However, in spiders, the type of eyes that undergo this modification varies depending on the ecological needs. For instance, jumping spiders of the family Salticidae have developed enlarged principal eyes to adapt to high-resolution vision in bright light conditions (Land, 1971). Conversely, spiders of the genus *Deinopis* have enlarged secondary eyes, rather than principal eyes, to adapt to nocturnal foraging (Stafstrom et al., 2017; Stafstrom & Hebets, 2016). Noticeably, some of the enriched terms in our positive selection analyses seem to be directly or indirectly related to eye size (*negative regulation of cell size* and *regulation of apoptotic signaling pathway*, *regulation of epithelial cell apoptotic process*, *negative regulation of endothelial cell apoptotic process*; Table S9 and Figures 6 and 7). Moreover, other significant terms are likely related to secondary eyes (*compound eye photoreceptor development* and *mushroom body development*; Table S9 and Figures 6 and 7). Since only secondary eyes direct visual information towards the mushroom bodies (Barth, 2002; Strausfeld & Barth, 1993), beneficial mutations causing selective sweeps in the candidate regions have been likely involved in adaptation to enhance *D silvatica* visual system in a similar way that in other species, probably to better detect movement in low-light-level environments.

Several species of the genus *Dysdera* exhibit preferentially, facultatively or even nearly obligate feeding on terrestrial woodlice. Species of this genus show adaptations for nutrient extraction, particularly with regard to nitrogen extraction efficiency (Toft & Macías-Hernández, 2017). Some components of the nitrogen metabolism and the urea cycle (*ornithine, citrulline, pyrroline−5−carboxylate, glutamate,* and *L-proline*) are among the significantly enriched GO terms in *D. silvatica* selective sweep candidate regions (Table S9 and Figures 6 and 7). In most animals, the urea cycle has a dual function: nitrogen excretion and arginine biosynthesis (Campbell, 1973). This amino acid is considered an essential nutritional and structural requirement (Rodriguez & Hampton, 1966), being involved in muscular contraction for providing rapid energy during fast movements (Laino et al., 2017) and the synthesis of proline, which is a predominant amino acid in cuticle proteins and in collagen (Cianciosi & Hird, 1986; Richards & Ireland, 1978). However, despite having a functional purine biosynthetic pathway, spiders and other arachnids have a non-functional or incomplete urea cycle, and excrete the excess of nitrogen as guanine instead of urea (Jenkinson et al., 1996). In these organisms, arginine becomes an essential dietary component (Campbell, 1973). Our results suggest that metabolic adaptations to increase nitrogen extraction efficiency from isopods (as a source of arginine) are still ongoing in this species. In fact, most of the metabolic differences observed between *Dysdera* with distinct dietary preferences can be attributed to changes in the performance of nitrogen extraction when fed on a different prey (Toft & Macías-Hernández, 2017), indicating that this trait is a major driver of species diversification in the Canary Islands. Besides, proline and arginine are also implicated in certain mechanical properties of silk. Proline can confer elasticity (Hayashi et al., 1999), while arginine may play a role in conferring resistance to excess humidity (Kim et al., 2021). Moreover, in scorpions, arginine metabolism can yield various products, some of which are involved in venom composition (Arjunwadkar & Reddy, 1983). Stimulatingly, a comparative transcriptomic study of five species of this genus identified two putative venom toxins (Vizueta et al., 2019) as candidates for adaptive changes in these species. Lastly, arginase, the enzyme responsible for converting arginine to proline and glutamate in nitrogen metabolism, uses manganese (Mn2+) as a cofactor. Interestingly, we found the biological process of ATP-dependent pumps transporting Mn2+ among the enriched GO terms in candidate selective sweep regions in our study.

Another noteworthy GO term enriched in candidate selective sweep regions is *chitin synthase activity* (Figure 7). Chitin, a complex biopolymer of sugar products, along with proteins, serves as the primary constituent of cuticles in arthropods. This term was also identified as a candidate by Vizueta et al., 2019. The genes associated with this function could be under positive selection to enhance protection or prevent desiccation in *D. silvatica*. We also observed the function of *calcium determination cytosol endoplasmic* among the significantly enriched terms. Calcium is one of the main intracellular signaling molecules, and it is involved in crucial processes such as muscle excitation-contraction and energy metabolism. Consequently, these functions are also potential targets of molecular adaptation to enhance prey performance in this species. Finally, based on our analysis, other functions could be under positive selection in *D. silvatica*, including fatty acid metabolism, glycometabolism and the production of energy and glycerol, which could be involved in rapid or seasonal adaptive responses to desiccation (preventing water loss), and cold stress (Coulson, 1990; Czajka & Lee, 1990; Danks, 2000; Misener et al., 2001; Williams et al., 2002). However, the candidate genes associated with the enriched GO terms related to these functions show orthologies with evolutionary distant genes or are difficult to interpret from a functional point of view. Although it is necessary to be cautious with the biological interpretations derived from all these results, we strongly contemplate previously introduced candidates being considered for future functional validation.

Overall, our study has revealed unexpected aspects of the recent evolution of an endemic island spider. The specific conditions favoring the large predicted ancestral effective population sizes and the recent population decline in this species are of special relevance to understand the origin, maintenance and loss of biodiversity in islands’ unique ecosystems, and therefore is instrumental to manage and conserve biodiversity in a changing world (Mergeay & Santamaria, 2012). On the other hand, selection analysis through the lens of population genomics points to traits such as vision and nitrogen extraction as targets of adaptation in this spider. The results presented here contribute to increasing our knowledge of a poorly studied arthropod lineage from the point of view of population genomics, in general and in particular, to have a clearer picture of how adaptive radiations arise in environments that are true natural laboratories of evolution.

## Acknowledgements

This work was supported by the Ministerio de Ciencia e Innovación of Spain (MCIN/AEI/10.13039/501100011033; grants PID2019-103947GB-C21 and PID2022-138477NB-C22 to J.R.; PID2019-105794GB and PID2022-137758NB-I00 to M.A.A; FPI fellowship BES-2017-081740 to P.E.), and from Comissió Interdepartamental de Recerca I Innovació Tecnològica of Catalonia, Spain (2021SGR00279). AS-G is a Serra Húnter fellow. We acknowledge the Garajonay National Parks for granting collection permits and helping with lodging and logistics during fieldwork.

## Availability of data

Raw sequence reads have been deposited at the DDBJ/ENA/GenBank under the Bioproject PRJNA1075441, with the following Sequence Read Archive (SRA) accession numbers, SRR27941309-SRR27941316 and SRR27941318-SRR27941321 for the *D. silvatica* low-coverage data, and the SRR28624427 and SRR27941317, the *D. silvatica* and *D. bandamae*, medium-coverage data.

## Supplementary Figure legends

**Supplementary Figure S1**. Phylogenetic tree of mitochondrial genomes. The scale bar represents 0.001 nucleotide substitutions per site.

**Supplementary Figure S2.** Mean per-site nucleotide diversity (*π*) and per-base per-generation recombination rate (*r*, estimated under the best-fit demographic model) in 50 kbp windows.

**Supplementary Figure S3.** Mean per-site nucleotide diversity (*π*) and per-base per-generation recombination rate (*r*, estimated constant size demographic model; see Material & Methods) in 5×10^-4^ bp windows for autosomes and the X-chromosome.

**Supplementary Figure S4.** Proportion of research articles published between 2010 and 2024 containing significant GO terms found in our study.

**Supplementary Figure S5.** Proportion of research articles published between 2010 and 2024 containing significant GO terms found in our study after the correction for multiple testing.

