## Supplementary figures and images for "Population genomics of adaptive radiations: Exceptionally high levels of genetic diversity and recombination in an endemic spider from the Canary Islands"

### Supplemental Figures

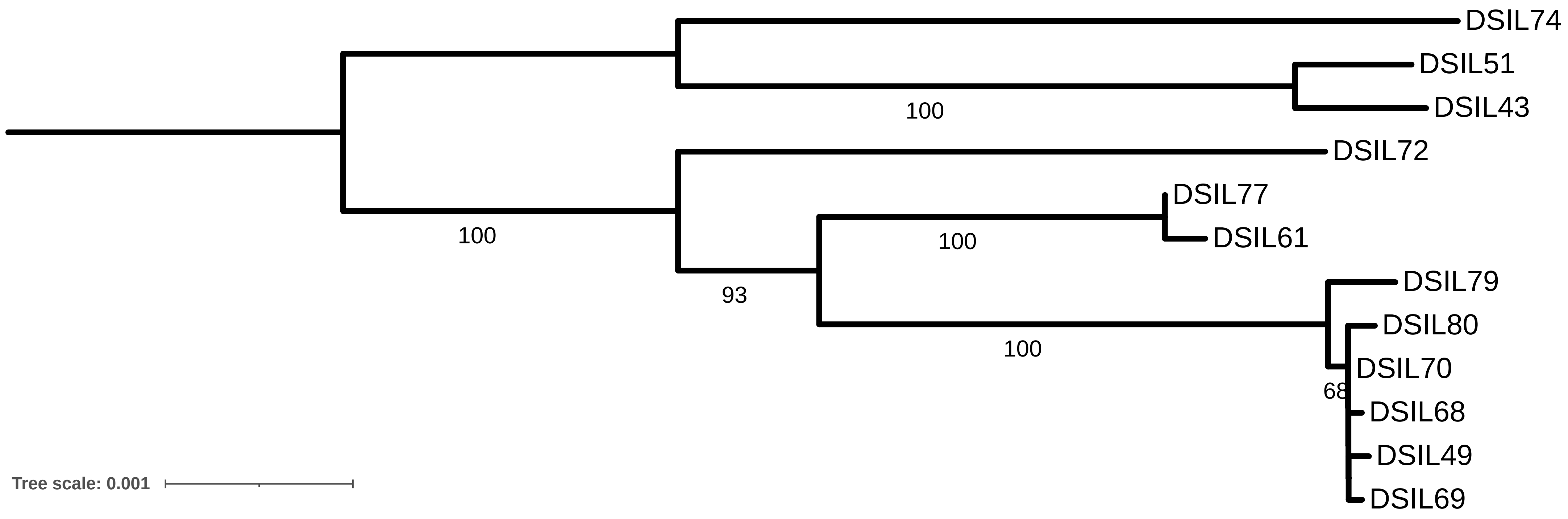

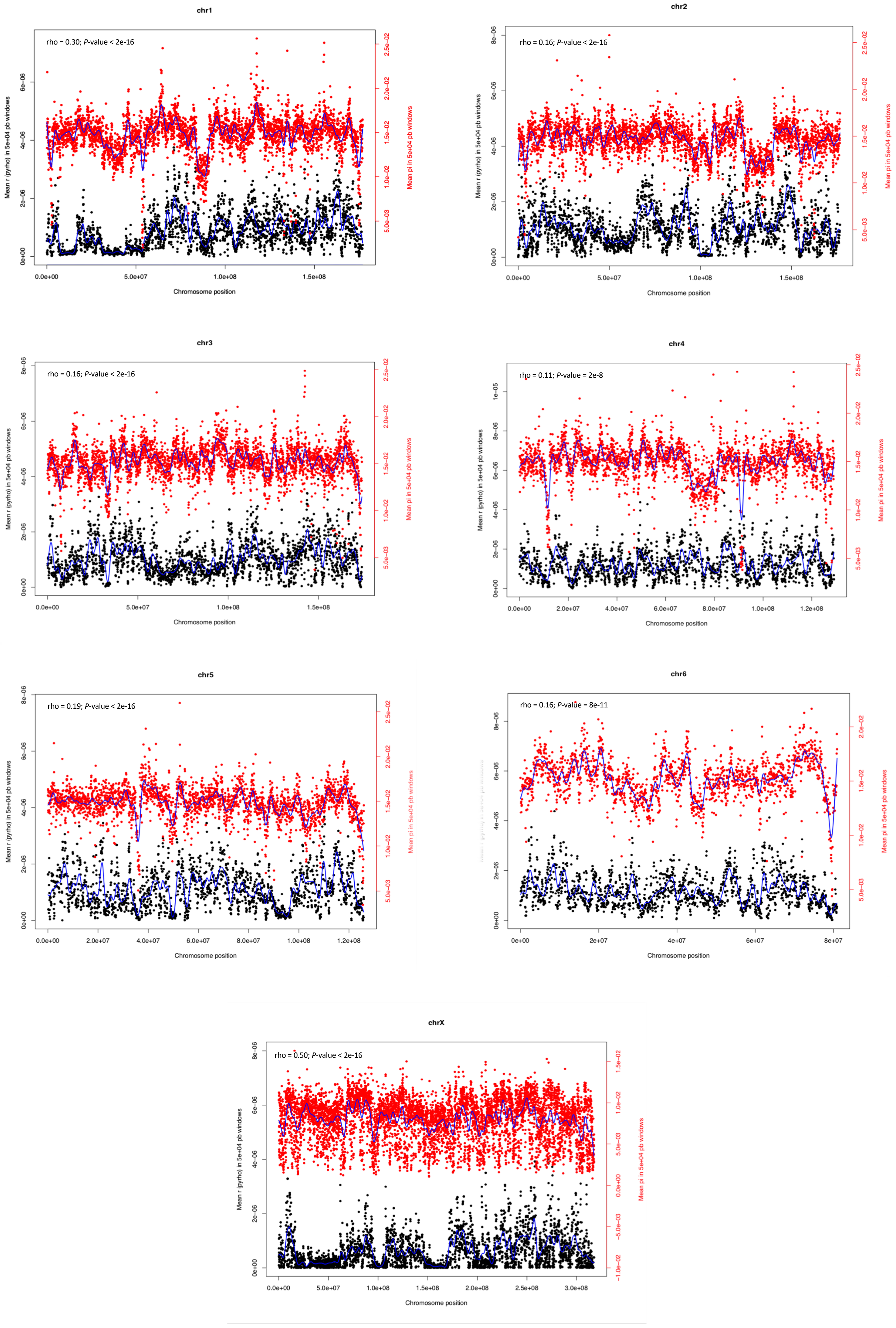

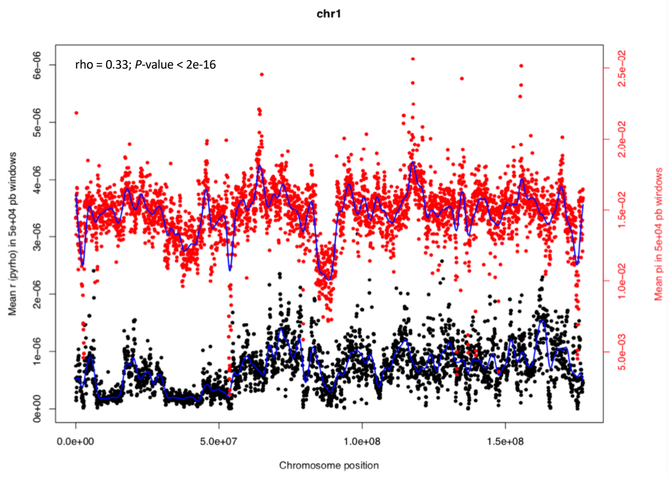

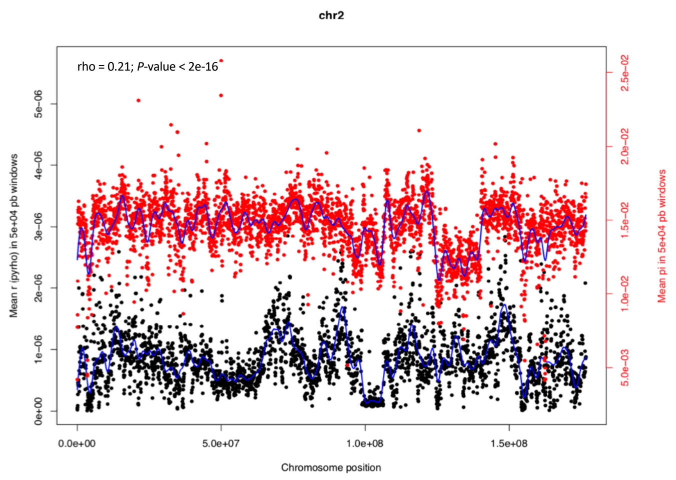

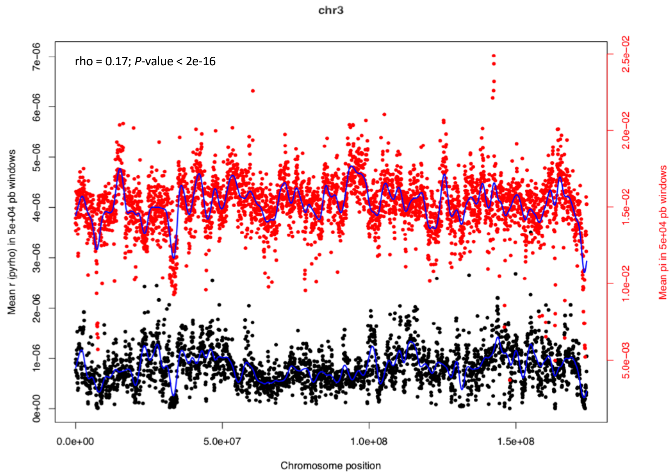

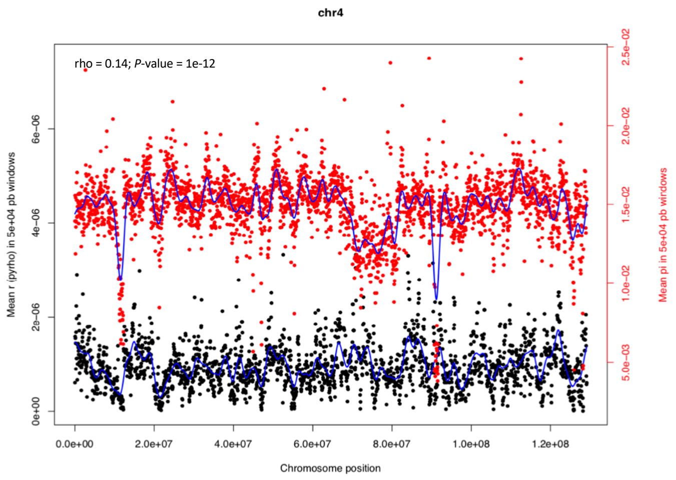

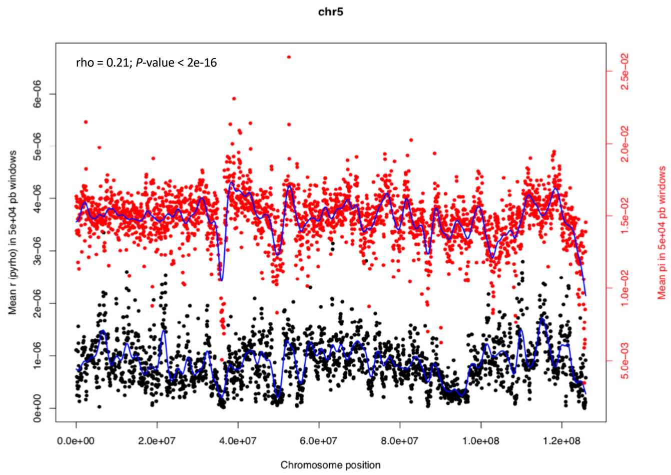

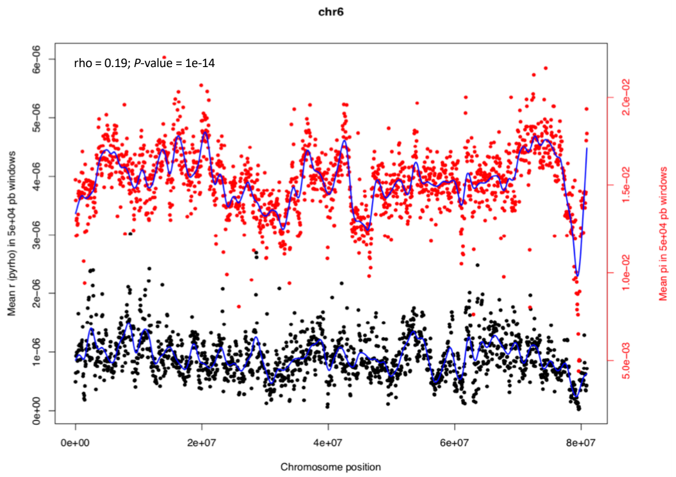

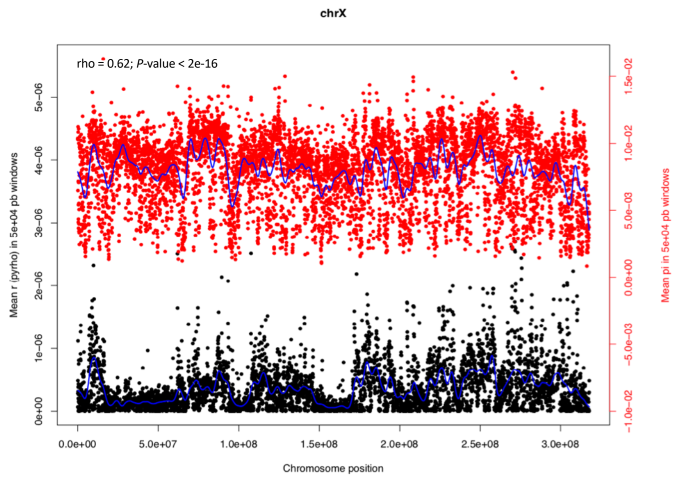

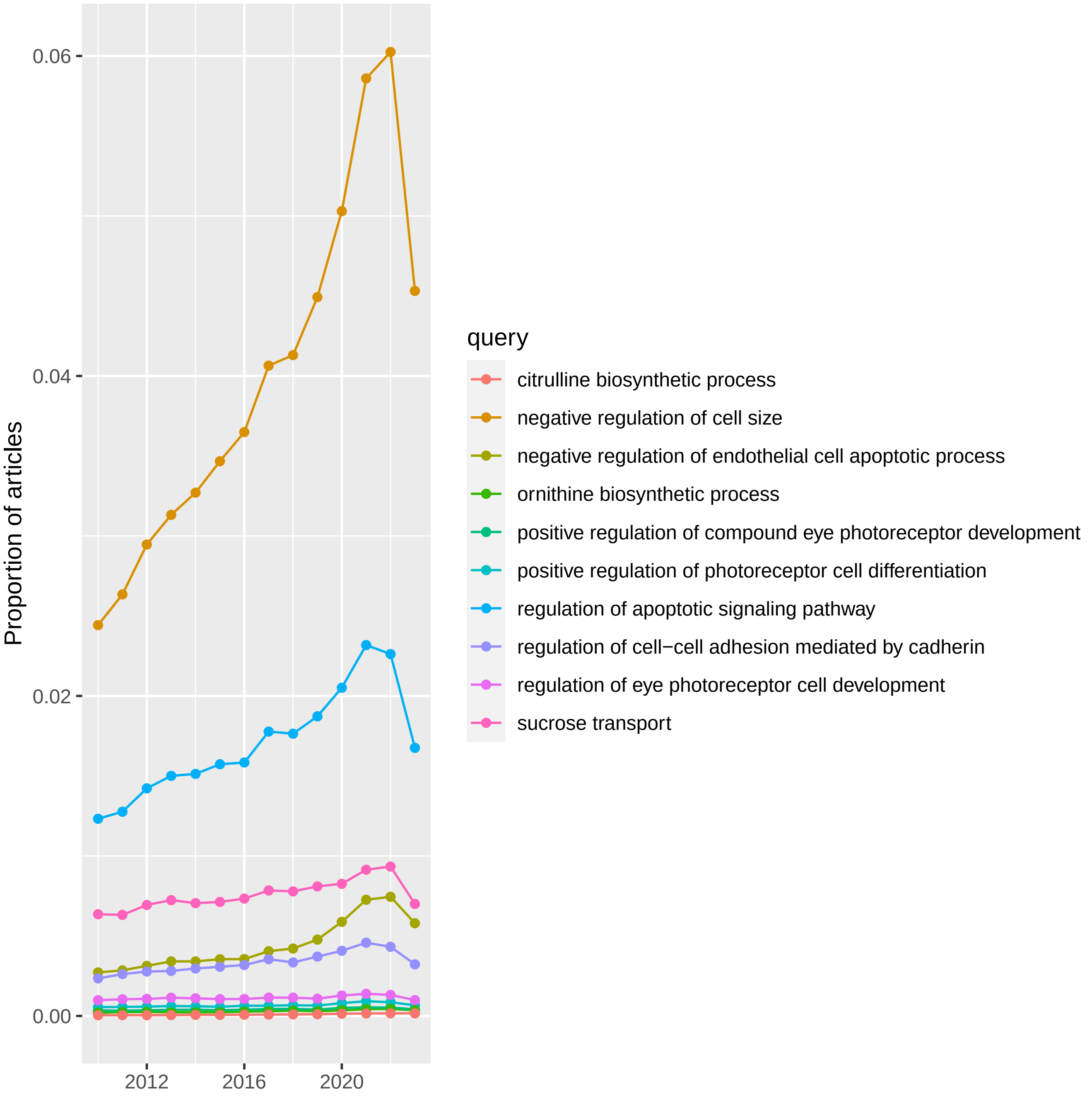

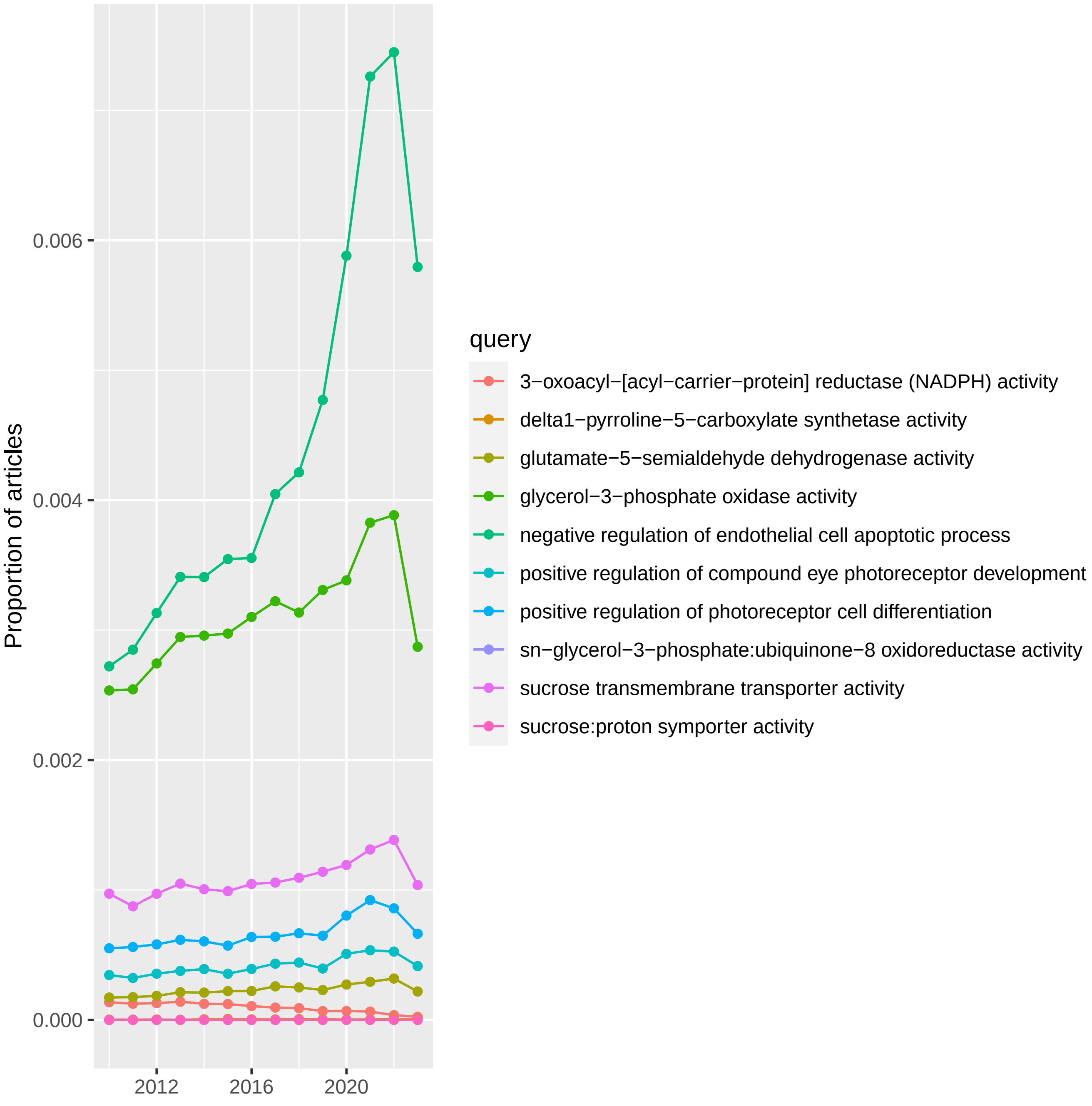
